# Integration of high-density genetic mapping with transcriptome analysis uncovers numerous agronomic QTL and reveals candidate genes for the control of tillering in sorghum

**DOI:** 10.1101/2020.07.06.189548

**Authors:** Rajanikanth Govindarajulu, Ashley N. Henderson, Yuguo Xiao, Srinivasa R. Chaluvadi, Margarita Mauro-Herrera, Muriel L. Siddoway, Clinton Whipple, Jeffrey L. Bennetzen, Katrien M. Devos, Andrew N. Doust, Jennifer S. Hawkins

## Abstract

Phenotypes such as branching, photoperiod sensitivity, and height were modified during plant domestication and crop improvement. Here, we perform quantitative trait locus (QTL) mapping of these and other agronomic traits in a recombinant inbred line (RIL) population derived from an interspecific cross between *Sorghum propinquum* and *Sorghum bicolor* inbred Tx7000. Using low-coverage Illumina sequencing and a bin-mapping approach, we generated ~1920 bin markers spanning ~875 cM. Phenotyping data were collected and analyzed from two field locations and one greenhouse experiment for six agronomic traits, thereby identifying a total of 30 QTL. Many of these QTL were penetrant across environments and co-mapped with major QTL identified in other studies. Other QTL uncovered new genomic regions associated with these traits, and some of these were environment-specific in their action. To further dissect the genetic underpinnings of tillering, we complemented QTL analysis with transcriptomics, identifying 6189 genes that were differentially expressed during tiller bud elongation. We identified genes such as Dormancy Associated Protein 1 (DRM1) in addition to various transcription factors that are differentially expressed in comparisons of dormant to elongating tiller buds and lie within tillering QTL, suggesting that these genes are key regulators of tiller elongation in sorghum. Our study demonstrates the usefulness of this RIL population in detecting domestication and improvement-associated genes in sorghum, thus providing a valuable resource for genetic investigation and improvement to the sorghum community.

## Introduction

Sorghum (*Sorghum bicolor* L. Moench), a staple cereal crop that grows in arid and semi-arid regions of the world, is an important food crop, a source of animal feed and forage, an emerging bioenergy crop (Rooney et al. 2007), and a model for C4 grasses, particularly maize, sugarcane, and *Miscanthus*. Among cereals, sorghum ranks fifth worldwide in acreage and annual tonnage after wheat, maize, rice and barley (Dogget 1998). Sorghum originated in Eastern Africa, but was taken by humans to Asia ~4000 years ago (Winchell et al. 2018). Varieties from Africa and Asia have subsequently contributed to the genetic makeup of current cultivated sorghum germplasm. The five races (bicolor, guinea, caudatum, kafir and durra) within *Sorghum bicolor* subsp. *bicolor* (2n = 2x = 20) are fully interfertile, and are derived from several wild progenitors, including *S. bicolor* subsp. *verticilliflorum*, *S. bicolor* subsp. *drummondii* and other weedy species that persist in sorghum growing regions of Africa (Dogget 1998; Mace et al. 2013; Wiersema and Dahlberg 2007). Several complex intermediate races have formed naturally or through breeding programs to achieve the most desired agronomic outcomes, such as increased yield, improved grain quality, and tolerance to various abiotic and biotic stressors (Mace and Jordan 2010, 2011; Morris et al. 2013). In addition, breeding efforts have produced sorghum germplasm that lacks photoperiod sensitivity, allowing its efficient introduction into temperate latitudes (Stephens et al. 1967).

Quantitative trait locus (QTL) mapping is a powerful approach to identify genomic regions that control specific phenotypes. Numerous sorghum recombinant inbred line (RIL) populations have been constructed, and QTL controlling >150 traits related to grain yield, leaf characteristics, maturity, panicle architecture, stem composition, stem morphology, perenniality, and abiotic and biotic stresses have been identified (reviewed in Mace and Jordan 2011). Nevertheless, the majority of the genetic diversity in wild and weedy crop relatives remains untapped, including in sorghum and related species from the center of diversity in Africa (Dogget 1995). To investigate the underlying genetic architecture controlling some of these traits, we generated an interspecific RIL population by crossing *S. propinquum* (female parent) with *S. bicolor* (Tx7000), representing the widest euploid cross that is easily performed in sorghum. Tx7000 is a variety known for pre-anthesis drought tolerance, and was used as an elite R-line for the production of early grain sorghum hybrids (Evans et al. 2013; Morishige et al. 2013). Prior crosses between Tx7000 and other inbred lines such as BTx642 (Evans et al. 2013), B35 (Subudhi et al. 2000; Xu et al. 2000) and SC56 (Kebede et al. 2001) have been used to investigate QTL related to drought tolerance and stay green traits. Tx7000 is most likely derived from Blackhull Kafir and Durra Dwarf Yellow Milo, and thus is substantially diverged from sorghum inbred BTx623, which was used to generate the sorghum reference genome sequence and as a parent in numerous segregating populations for the construction of genetic maps (Evans et al. 2013; Mace and Jordan 2011). *S. propinquum* is a wild relative, indigenous to SE Asia, that has small seeds, high density tillering, narrow leaves, day-length-dependent flowering, and well-developed rhizomes that contribute to perenniality. Because of the many phenotypic differences between *S. bicolor* and *S. propinquum*, the RIL population developed in this study provides an opportunity to investigate agronomically important traits such as tillering and aerial branching, plant height, flowering time, and biomass, all of which are important in grain, forage, and fuel production (Jessup et al. 2017; Rooney et al. 2007).

QTL mapping can be combined with transcriptome analysis (RNA-seq) to identify candidate genes underlying QTL for traits of interest. For example, Gelli et al. (2014) identified candidate genes associated with nitrogen stress tolerance in sorghum by overlaying differentially expressed genes onto validated QTL between parents and pools of RILs with high and low nitrogen use efficiency (NUE). Similarly, the genetic basis of O_3_ sensitivity was examined in Arabidopsis by the combined application of QTL mapping and differential transcriptome analysis of the parents (Xu et al. 2015). Here, we evaluated several agronomically important traits in our RILs at two field sites and in the greenhouse. We complemented the QTL mapping of tillering with transcriptome analysis during tiller bud elongation. These experiments enabled the identification of 30 QTL related to six important agronomic traits in addition to several candidate genes associated with tiller bud dormancy and elongation.

## Materials and methods

### Plant material

We developed a mapping population consisting of 191 F_3:5_ RILs derived from a cross between *Sorghum propinquum* (female, unnamed accession obtained from Dr. William Rooney, Texas A&M University, College Station, TX) and *S. bicolor* inbred “Tx7000”. Each RIL was derived from a single F_2_ plant following the single seed descent method and progressed up to the F_5_ generation. Because of day length issues for flowering at some locations, some RILs trailed the overall population advancement, and are now only at the F_3_ or F_4_ generations. For the transcriptome analysis, we germinated seeds and collected tiller buds for mRNA sequencing. We used Tx7000 and *S. bicolor* subsp. *verticilliflorum* (PI302267) for this part of the study because seed germination for *S. propinquum* was low and asynchronous with Tx7000, preventing informative comparisons at 8 and 14 days post-planting. *S. bicolor* subsp. *verticilliflorum* produces tillers and is phylogenetically closer to *S. propinquum* than any other cultivated or wild species of sorghum (Mace et al. 2013).

### Field trial design and greenhouse study

Phenotypic performance of the RILs and parents was evaluated at two field locations and in one greenhouse experiment (**Table 1**). Field trials were conducted at the University of Georgia’s Iron Horse Farm, Watkinsville, GA, USA (33.725° N, 83.3° W; coarse sandy loam soil) and at the Oklahoma State University’s Cimarron Valley Agricultural Research Station, Stillwater, OK, USA (35.986° N, 97.049° W; sandy loam soil) in the years 2016 and 2017, respectively. Henceforth, these populations will be referred to as 2016-W and 2017-S. Seeds were germinated in mid-April for both field studies. In 2016-W, fifteen plants from each RIL were transplanted in mid-May in a randomized complete block design with three replications. Five seedlings per RIL were planted in each replication in a row, with 61 cm between rows and 51 cm between plants within a row. In 2017-S, 30 plants from each RIL were transplanted in a randomized complete block design with three replications and containing two replications per block. Spacing was the same as that of the 2016-W experiment. Standard agronomic practices were followed throughout the growing season, with supplemental irrigation and weed control as necessary.

**Table 1.** Summary of sorghum Recombinant Inbred Lines (RILs) at each study site. Number of RILs from each seed generation grown at each study location are shown. For each RIL, only the most advanced generation was used.

| Generation of RIL |  |  |  | Number of RILs |
| --- | --- | --- | --- | --- |
| Location | F3 | F4 | F5 |  |
| 2014 - GH | 9 | 42 | 120 | 171 |
| 2016 - W | 17 | 46 | 119 | 182 |
| 2017 - S | 13 | 42 | 115 | 170 |

**Locations:** 2014 – GH, Greenhouse, Morgantown, WV; 2016 – W, Watkinsville, GA; 2017 – S, Stillwater, OK

The greenhouse experiment was conducted in July 2014, and will henceforth be referred to as 2014-GH (**Table 1**). Nine seeds from each RIL (3 replicates with 3 plants per line per replication) were planted in germination mix (Metro-mix, Sun Gro Horticulture, Bellevue, WA, USA), and transplanted two weeks after germination into 15 cm Azalea pots (1.3 liter capacity) filled with Metro-mix soil (Sun Gro Horticulture, Bellevue, WA, USA). Plants were grown in small pots to evaluate their ability to tiller in limited growing space. Plants were grown at 21°C/26°C night/day temperatures under natural day length. Plants were fertilized once every three weeks with 20:18:20 N:P:K (Jacks water-soluble fertilizer, Allentown, PA) in addition to 40 grams of granular slow-release Osmocote 14:14:14 N:P:K fertilizer (Osmocote Pro). Plants were treated with 15 grams of Ironite (Gro Tec, Inc, Madison, GA) as needed.

### Genotype analysis using whole genome re-sequencing

High-quality nuclear DNA was isolated from the parents and the RILs using a nuclear isolation protocol (Lutz et al. 2011; Peterson et al. 1997), specifically Protocol B as described by Lutz and coworkers, which maximizes nuclear genome content by limiting organellar contamination. Illumina sequencing libraries with dual-indexes were prepared following the method of Glenn et al. (2016). The 100 bp paired-end (PE) sequencing was performed on an Illumina HiSeq 2000 at either the Marshall University Genomics Core Facility, Huntington, WV or the University of Georgia Genomics and Bioinformatics Core, Athens, GA. Parental lines and RILs were sequenced at ~18x and ~2x depths, respectively. For SNP calling, the sequence reads from each RIL were aligned to the masked *S. bicolor* reference, ver 3.1 (Paterson et al. 2009) using Bowtie2 (Langmead and Salzberg 2012) with default parameters. SNPs were called using samtools mpileup (Li et al. 2009). High quality SNPs with 1) a minimum genotype quality score of 30, 2) a minimum sequencing depth of 2 for each RIL, and 3) called in at least 40% of the RILs were selected for the construction of genetic maps for subsequent QTL analysis.

### Genetic map construction

The high-quality SNP variant call file was parsed through the GenosToABHPlugin in Tassel ver 5.0 (Bradbury et al. 2007) to obtain informative biallelic SNPs in a parent-based format (*S. propinquum* alleles as “A”, *S. bicolor* alleles as “B”, and heterozygotes as “H”), and to exclude sites in which the parental genotypes were missing, ambiguous, or heterozygous. The ABH-formatted SNP data file was used as input to SNPbinner (Gonda et al. 2019), which calculates breakpoints (transition between haplotypes) in each RIL and constructs genotype bins that are implemented in three python modules (crosspoints.py, visualize.py, and bins.py). Briefly, the module crosspoints.py, which uses a pair of hidden Markov models (HMM), one considering all three states (A, B and H) and the other considering homozygous states (A and B), was used to determine the most likely genotype at each point along a chromosome for each RIL. Breakpoints that were separated by less than 0.2% of the chromosome length were merged into the surrounding regions. The visualize.py module was used on a random selection of chromosomes to visualize the representation of the original SNPs against the calculated breakpoints for quality control and to determine the optimal parameters for crosspoint calculations. The bins.py module was used to create a combined map of all breakpoints across the RILs at a user-specified resolution (a 10 kb bin size was used in this study). Finally, duplicate bin markers and markers that were potentially indicative of double-crossovers (as determined by calculating genotyping error LOD scores using calc.errorlod() function in R/qtl) were removed, and the remaining markers were used to construct the genetic map. The markers were ordered based on the physical positions of the bins in the reference genome, and the centimorgan distances were calculated using the Kosambi map function in R/qtl (Broman et al. 2003) (**Table 2**).

**Table 2.** Construction of genetic maps using bin mapping. Physical chromosome length, total number of SNPs considered per bin, total bin markers used in R/qtl, number of bin markers after removing co-segregating markers, average bin size, genetic map distance, and number of break points per chromosome are shown.

| Chr | Chr length (Mb) | Total number of SNPs post filtering |  |  | Number of bin markers on the map |  |  | Average bin size (Mb) |  |  | Genetic map distance (cM) |  |  | Number of breakpoints |  |  |
| --- | --- | --- | --- | --- | --- | --- | --- | --- | --- | --- | --- | --- | --- | --- | --- | --- |
|  |  | 2014 | 2016 | 2017 | 2014 | 2016 | 2017 | 2014 | 2016 | 2017 | 2014 | 2016 | 2017 | 2014 | 2016 | 2017 |
| 1 | 80.9 | 61534 | 29306 | 48390 | 256 | 236 | 237 | 0.14 | 0.15 | 0.15 | 113.4 | 106.8 | 112.5 | 913 | 935 | 880 |
| 2 | 77.7 | 41335 | 17254 | 29543 | 239 | 234 | 239 | 0.17 | 0.18 | 0.17 | 98.6 | 93.2 | 100.1 | 769 | 788 | 744 |
| 3 | 74.4 | 49257 | 21032 | 35198 | 222 | 230 | 219 | 0.17 | 0.17 | 0.18 | 124.0 | 121.8 | 124.8 | 823 | 874 | 796 |
| 4 | 68.7 | 42096 | 18169 | 30178 | 198 | 200 | 198 | 0.17 | 0.18 | 0.17 | 86.9 | 82.6 | 83.4 | 748 | 761 | 712 |
| 5 | 71.8 | 21359 | 9415 | 15398 | 159 | 161 | 155 | 0.24 | 0.24 | 0.25 | 77.1 | 73.2 | 76.2 | 652 | 689 | 622 |
| 6 | 61.3 | 35065 | 15321 | 25299 | 154 | 148 | 147 | 0.18 | 0.19 | 0.20 | 65.2 | 60.1 | 64.8 | 652 | 645 | 616 |
| 7 | 65.5 | 27829 | 11939 | 20138 | 171 | 179 | 177 | 0.19 | 0.20 | 0.20 | 82.6 | 79.7 | 83.4 | 669 | 712 | 661 |
| 8 | 62.7 | 22320 | 9660 | 15520 | 163 | 158 | 155 | 0.23 | 0.24 | 0.24 | 78.3 | 71.7 | 76.6 | 576 | 607 | 569 |
| 9 | 59.4 | 30546 | 12659 | 21276 | 201 | 194 | 196 | 0.17 | 0.17 | 0.17 | 87.4 | 80.3 | 88.7 | 710 | 730 | 680 |
| 10 | 61.2 | 28777 | 12207 | 20508 | 171 | 175 | 183 | 0.18 | 0.19 | 0.19 | 77.5 | 77.2 | 78.7 | 674 | 698 | 648 |
| **Total** | **683.5** | **360118** | **156962** | **261448** | **1934** | **1915** | **1906** | **0.18** | **0.19** | **0.19** | **891.2** | **846.6** | **889.2** | **7186** | **7439** | **6928** |
| **Total bin markers** |  |  |  |  | **3803** | **3666** | **3652** |  |  |  |  |  |  | **4.20** | **4.09** | **4.08** |

### Phenotype evaluation

Three individual plants from the middle of the five consecutive plants (plants 2, 3 and 4) in each of the three replicate blocks were scored for phenotypes in 2016-W (total 9 plants), while two plants from the middle of the five consecutive plants (plants 2 and 4) in each of the 6 replicates were scored for phenotypes in 2017-S (total of 12 plants). At maturity, plant height (HT) was measured from ground to the apex (panicle tip) if flowering, and if the plant was not flowering, height was measured from the ground to the base of the top leaf. Tiller number (TN) was recorded by counting the number of tillers that had emerged from the base of the plant. Stem diameter (SD) was measured using vernier calipers between the first and second aboveground internode. The days to flowering (FL) was determined by collecting the heading dates for each plant in the greenhouse trial and in the 2017-S field trial. FL was not recorded during the 2016-W field experiment. To obtain dry weight for biomass (B) measurements, the aboveground portion of the plant was manually harvested from both field trials, placed in separate Mesh harvest bags (Midco, USA), and placed in a drying oven (140 °F) for one week. Branches (ARB) emerging from the internodes were counted during the 2017-S field experiment.

### Phenotype analysis

The total number of RILs evaluated in each experiment is indicated in **Table 1**. Least square means for each trait were calculated for each environment across replicates and examined for normality using Q-Q plots in R (R core team 2013). Traits that deviated from normality were transformed as necessary, and the transformed data were used in an analysis of variance (ANOVA). In the greenhouse experiment, flowering date and stem diameter were transformed using Tukey ladder of powers transformation with an appropriate lambda value (Tukey 1977). For the 2017-S field experiment, tiller number and height were square-root transformed. For the 2016-W field experiment, log transformation was applied to biomass and box-cox transformation was applied to height.

Correlation of phenotypic traits within each site was explored using Pearson correlation in R (R core team, 2013) (**Supplemental Figure S1**). Combined analysis of variance was performed by fitting a linear model (lm) to assess genotype (G), environment (E), and genotype x environment (GxE) interactions in R (lm and calculated Type III analysis of variance (ANOVA) summary for the model) (**Supplemental Table S1**). Because of the significant GxE interaction, we generated the best linear unbiased predictors (BLUP) for three traits (tiller number, height, and stem diameter) that were measured similarly in both field environments. To generate BLUPs, block and location were considered as fixed effects, whereas G and GxE were considered as random effects. The BLUP values were used in QTL analysis.

### QTL analysis

QTL analysis was performed in R/qtl ver.1.14 (Broman et al. 2003) using only the RILs that were both genotyped and phenotyped. For QTL analyses, we used genetic maps that were constructed using 1934, 1915, and 1906 bin markers from the 2014-GH, 2016-W, and 2017-S experiments, respectively (**Table 2**). The genetic maps consisted of 10 linkage groups that correspond to the ten chromosomes of the sorghum genome (Paterson et al. 2009). QTL mapping was performed for each field experiment separately. To summarize phenotypic values for correlated traits (tiller number, height, and stem diameter) from the two field sites, we used 170 common RILs that yielded 1906 bin markers for the BLUP analysis. A single QTL model using an interval mapping method and appropriate phenotype model was chosen to identify QTL exhibiting logarithm of the odds (LOD) score peaks greater than the permutation-based significance threshold (alpha = 0.05) estimated via 1000 permutations (Churchill and Doerge 1994). Next, a multiple QTL model that uses forward selection and backward elimination to determine the best fit QTL model was used to identify additional additive QTL, refine the QTL positions, and test for the interaction between QTL for a given trait. The significance of fit for the full multiple QTL model was assessed using Type III ANOVA. The proportion of variance explained and the additive effect for each locus was extracted from the model summary. Because of the large window size obtained using MQM for tillering QTL, composite interval mapping (CIM) was implemented in R/qtl using a genome scan interval of 1 cM, a window size of 10 and a forward and backward regression method to refine the QTL position (Jansen and Stam 1994; Zeng 1994). Putative genes (*Sorghum bicolor* ver 3.1) found within a 1.0 LOD confidence interval for each QTL were identified.

### RNA isolation, RNA-seq library construction and sequencing

*S. bicolor* inbred Tx7000 and *S. bicolor* subsp. *verticilliflorum* plants used for RNA-seq were grown in controlled growth chambers at 24°C, with a 16-h light/8-h dark cycle. Tiller buds collected at 10, 12, and 14 DAP (days after planting) were dissected from the first leaf axil under a stereomicroscope and flash frozen in liquid nitrogen. Bud size measurements were taken at 8, 10, 12, 14 and 16 DAP to compare early tiller bud development between the two genotypes. All sampling was performed within a two-hour window in the afternoon approximately 5-7 hours after dawn to minimize circadian effects. Depending on the size of tiller buds at a given developmental stage, 50–144 individual tiller buds were pooled per biological replicate, and three biological replicates were collected per sample. RNA was isolated, for three replicates, from tiller buds collected at 10, 12, and 14 DAP, using the QIAGEN miRNeasy Micro kit (cat. no. 217084) according to the manufacturer’s instructions. The RNA-seq libraries were prepared using a KAPA stranded mRNA kit (cat.no. KK8421) according to the manufacturer’s instructions (KAPABIOSYSTEMS, https://www.kapabiosystems.com). The average library insert size was approximately 500 bp. Libraries were assessed for quality and quantified on an Agilent bioanalyzer (Agilent), and 125 bp paired-end Illumina (Hi-seq 2500) sequencing was performed at the Marshall University Genomics Core Facility (Marshall University, Huntington, WV).

### Differential expression analysis of RNA-seq data

Methods for differential expression analysis were as described in Dong et al. 2019. Eighteen RNA-seq libraries were sequenced and ~428 million paired-end (PE) raw reads were obtained, averaging ~23 million reads per library. Overall quality was assessed using FastQC (Andrew 2010). Trimmomatic v.0.36 (Bolger et al. 2014) was used to remove Illumina adaptor sequences, trim bases with quality scores less than 30 from the 5’ and 3’ end of each read, and to discard reads of less than 36 bp in length. The processed reads were mapped to the reference genome, version 3.1 (Sb.ver3.1), of *S. bicolor* BTx623 using STAR aligner v.2.6.0a (Dobin et al. 2012) with default settings. The total and uniquely mapped reads are summarized in **Supplemental Table S2**. A read count matrix was generated by aggregating the raw counts of the mapped reads for a given gene in each sample using featureCounts (Liao et al. 2013) in reference to 34,211 sorghum gene models in Sb.ver3.1 (**Supplemental Table S3**). The read count matrix was then employed for differential gene expression analysis using the R package edgeR v.3.22.5 (Robinson et al. 2010). In order to improve sensitivity, only genes with a count-per-million (CPM) value > 1 in at least three libraries were used for gene expression analysis. The filtered read count matrix was normalized for compositional bias between libraries using the trimmed means of M values (TMM) method, and differentially expressed genes were identified from pairwise comparisons of samples. Genes with an adjusted p-value (q-value) ≤ 0.05 and an absolute value of log2 fold changes (FC) ≥ 1 were considered differentially expressed. We identified the differentially expressed genes that lie within 1 LOD of the physical position of the tillering QTL peak in the *S. bicolor* reference genome.

### Co-expression cluster analysis and GO enrichment analysis

Differentially expressed genes (DEGs) (**Supplemental Table S4**) were subjected to co-expression cluster analysis, both among species and between developmental stages, using the R package coseq v1.5.2 (Rau and Maugis-Rabusseau 2017). The RPKM (Reads Per Kilobase of transcript per Million mapped reads) values for all DEGs were extracted and used as input for co-expression cluster analysis. Briefly, log CLR-transformation and TMM normalization were applied to the gene expression matrix to normalize the expression of genes, and the K-means algorithm was chosen to detect the co-expressed clusters across all samples. The K-means algorithm was repeated 30 times in order to determine the optimal number of clusters. The resulting number of clusters in each run was recorded, and then the most parsimonious cluster partition was selected using an adjusted random index (compareARI function in coseq). Statistically enriched (adjusted p-value ≤ 0.05) gene ontology (GO) terms for differentially expressed genes between pairwise comparisons, or of genes assigned to each co-expression cluster, were identified using Singular Enrichment Analysis (SEA) in AgriGO v2.0 with the *S. bicolor* v2.1 genome sequence as a background (Tian et al. 2017). The most statistically enriched GO terms were plotted in ggplot2 (Wickham 2016) for visualization.

## Results

### Construction of genetic maps using bin mapping

After quality filtering, 360,118, 156,962, and 261,448 SNPs were retained for bin mapping in the 2014-GH, 2016-W, and 2017-S populations, respectively (**Table 2**). In the RILs assessed in the 2016-W and 2017-S field trials, we detected 7439 and 6928 breakpoints from 182 and 170 RILs with an average of 4.09 and 4.08 breakpoints per chromosome per RIL, respectively. In the greenhouse experiment, there were 7186 breakpoints in 171 RILs with an average of 4.20 breakpoints per chromosome per RIL. A total of 3803, 3666, and 3652 bin markers were obtained from grouping the filtered SNPs sites for the 2014-GH, 2016-W, and 2017-S field experiments, respectively. After removing duplicate markers and double crossovers, the genetic maps included 1934, 1915, and 1906 bin markers from the 2014-GH, 2016-W, and 2017-S experiments, respectively, and the average physical length of the bins per chromosome was ~180 kb, ranging from 10 kb to 17 Mb (**Table 2**). The total genetic distance of the bin map across the three trials averaged 875 cM, with ~3.6 cM per bin. The average retained heterozygosity, a measurement based on the proportion of heterozygous bin markers, was 16%. The recombination bin map for the common 170 RILs from the 2016-W and 2017-S experiments is shown in **Supplemental Figure S2**.

### Segregation distortion

The physical distribution of segregation distorted bin markers was visualized by plotting the physical bin marker positions versus the associated p-value (converted in terms of negative logarithm with a base of 10) for the 170 common RILs from the 2016-W and 2017-S field experiments (**Supplemental Figure S3**). A chi-squared test was used to calculate the deviation from the expected Mendelian ratio (1:1) for each bin marker. Distorted markers (deviation at 0.1% [p< 0.001]) were distributed across eight of the ten chromosomes. The distorted regions on chromosomes 2 (peak at 65 Mb) and 5 (peak at 12.9 Mb) are enriched for *S. propinquum* alleles, whereas the highly distorted regions on chromosomes 1 (peak at 62.5 Mb), 4 (peak at 66.1 Mb), 6 (peak at 0.7 Mb), and 10 (peak at 60 Mb) are enriched for *S. bicolor* alleles (**Table 3**).

**Table 3.**
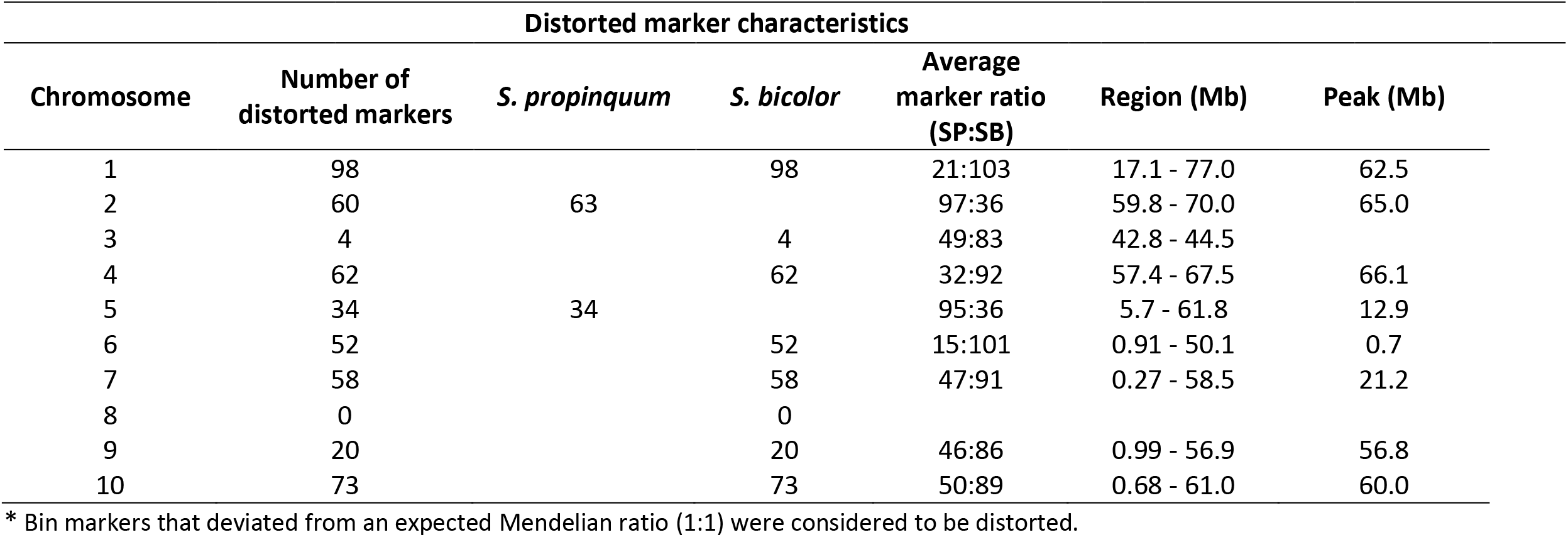
Bin markers with highly distorted segregation ratios on eight of the ten chromosomes.

### Phenotype and QTL analysis using the high-density bin map

#### Tillering

*S. propinquum* produced a maximum of 16 tillers, while Tx7000 produced zero to four tillers, in the field. (**Table 4**). In the greenhouse, *S. propinquum* produced 1 or 2 tillers, but Tx7000 produced no tillers. The average tiller number in the RIL population in the field trials was 1 (2016-W) and 5 (2017-S), with a maximum of 27. In all three experiments, tillering was positively skewed (skewness ≥ 1) (**Supplemental Figure S4a**). Two QTL for tillering on chromosomes 1 and 7 were detected in both field trials (**Table 5**). BLUPs summary of tillering validated the QTL on chromosomes 1 and 7 with peaks at 15.86 Mb and 8.36 Mb (**Figure 1)**, respectively, that together explained 31 percent of the phenotypic variation. These QTL (qTN1.blup and qTN7.blup) had an average negative additive effect of −0.54, indicating that the Tx7000 alleles negatively influence tillering. In the greenhouse, tillering QTL (qTN3.14GH, qTN9.14GH) were detected on chromosomes 3 and 9 with peaks at 62.01 Mb and 45.79 Mb that together explained 31 percent of the variation.

**Table 4.**
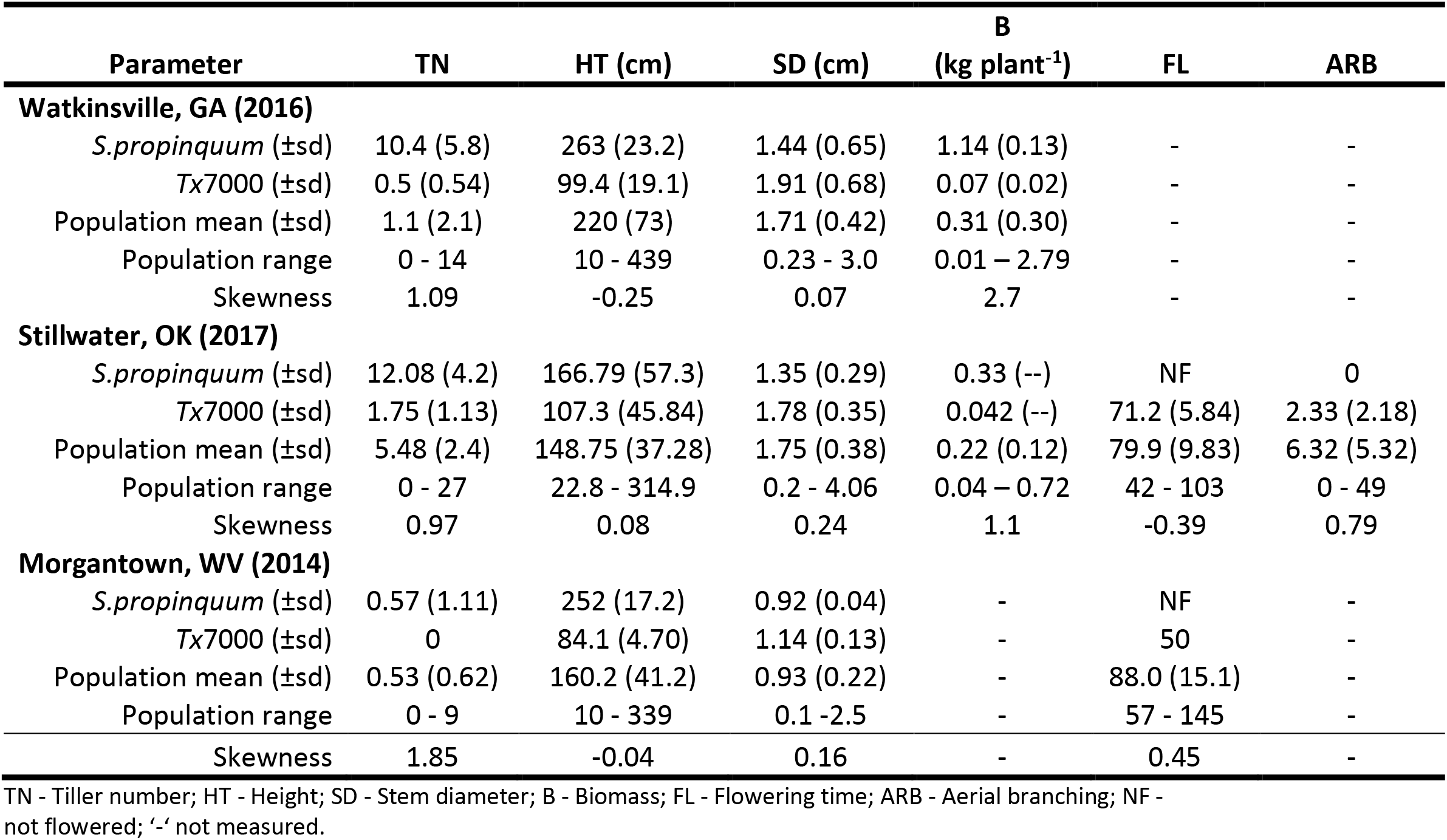
Summary of the trait data for the RIL mapping population compared to parental performance in Watkinsville, GA (2016), Stillwater, OK (2017) and in the Greenhouse, Morgantown, WV (2014).

| Parameter | TN | HT (cm) | SD (cm) | B<br>(kg plant<sup>-1</sup>) | FL | ARB |
| --- | --- | --- | --- | --- | --- | --- |
| **Watkinsville, GA (2016)** |  |  |  |  |  |  |
| *S.propinquum* (±sd) | 10.4 (5.8) | 263 (23.2) | 1.44 (0.65) | 1.14 (0.13) | - | - |
| *Tx7000* (±sd) | 0.5 (0.54) | 99.4 (19.1) | 1.91 (0.68) | 0.07 (0.02) | - | - |
| Population mean (±sd) | 1.1 (2.1) | 220 (73) | 1.71 (0.42) | 0.31 (0.30) | - | - |
| Population range | 0 - 14 | 10 - 439 | 0.23 - 3.0 | 0.01 – 2.79 | - | - |
| Skewness | 1.09 | -0.25 | 0.07 | 2.7 | - | - |
| **Stillwater, OK (2017)** |  |  |  |  |  |  |
| *S.propinquum* (±sd) | 12.08 (4.2) | 166.79 (57.3) | 1.35 (0.29) | 0.33 (--) | NF | 0 |
| *Tx7000* (±sd) | 1.75 (1.13) | 107.3 (45.84) | 1.78 (0.35) | 0.042 (--) | 71.2 (5.84) | 2.33 (2.18) |
| Population mean (±sd) | 5.48 (2.4) | 148.75 (37.28) | 1.75 (0.38) | 0.22 (0.12) | 79.9 (9.83) | 6.32 (5.32) |
| Population range | 0 - 27 | 22.8 - 314.9 | 0.2 - 4.06 | 0.04 – 0.72 | 42 - 103 | 0 - 49 |
| Skewness | 0.97 | 0.08 | 0.24 | 1.1 | -0.39 | 0.79 |
| **Morgantown, WV (2014)** |  |  |  |  |  |  |
| *S.propinquum* (±sd) | 0.57 (1.11) | 252 (17.2) | 0.92 (0.04) | - | NF | - |
| *Tx7000* (±sd) | 0 | 84.1 (4.70) | 1.14 (0.13) | - | 50 | - |
| Population mean (±sd) | 0.53 (0.62) | 160.2 (41.2) | 0.93 (0.22) | - | 88.0 (15.1) | - |
| Population range | 0 - 9 | 10 - 339 | 0.1 - 2.5 | - | 57 - 145 | - |
| Skewness | 1.85 | -0.04 | 0.16 | - | 0.45 | - |

TN - Tiller number; HT - Height; SD - Stem diameter; B - Biomass; FL - Flowering time; ARB - Aerial branching; NF - not flowered; '-' not measured.

**Table 5.**
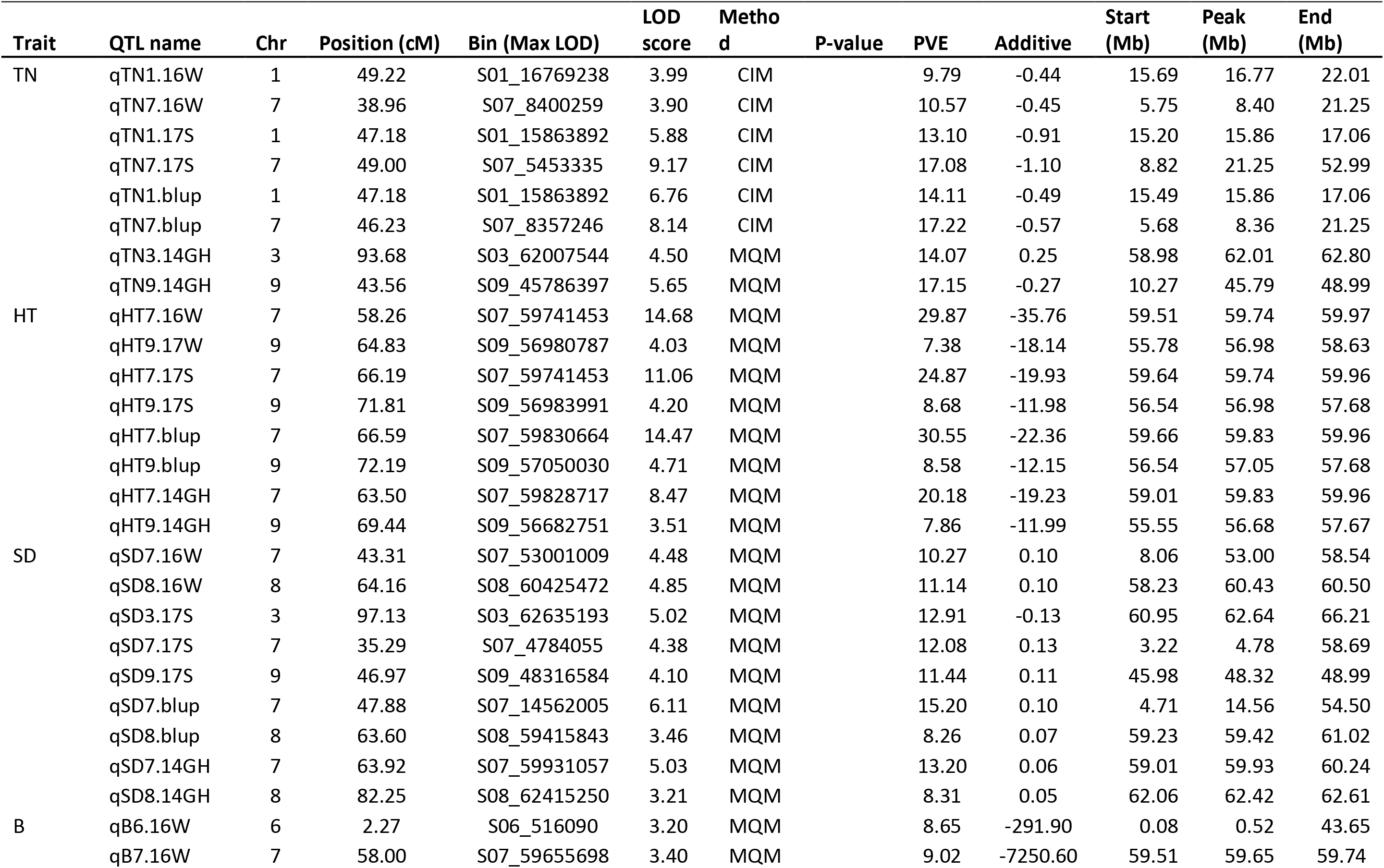
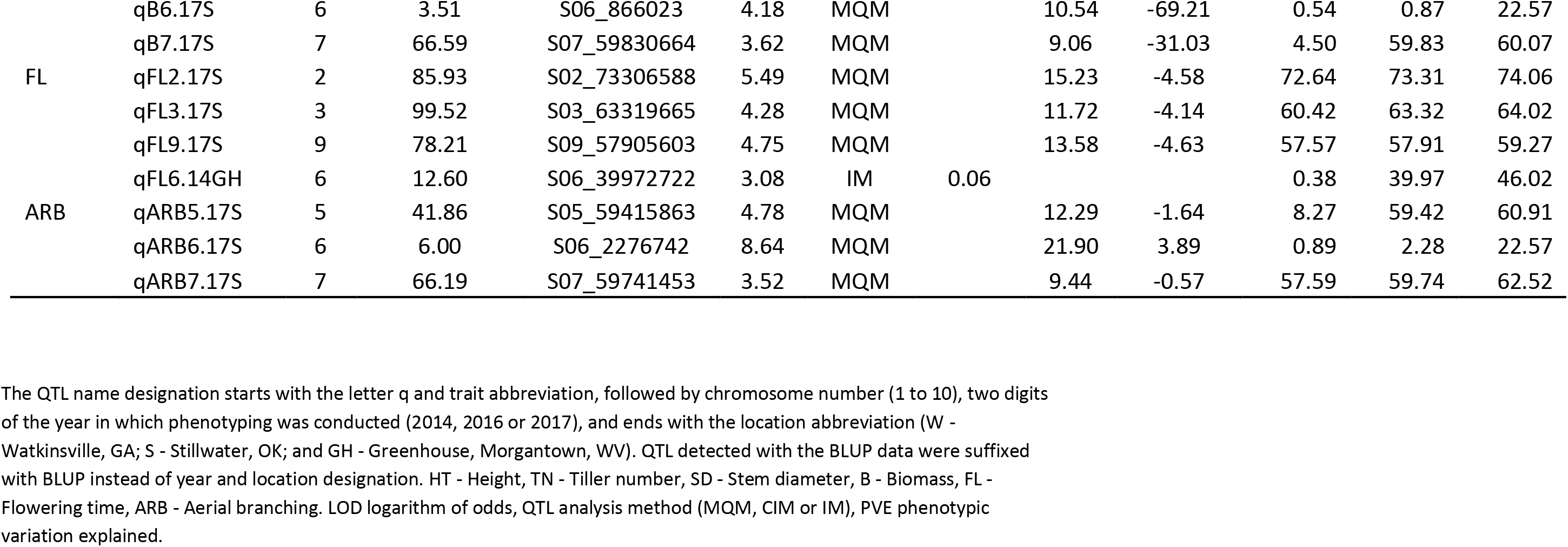
QTL identified in the RIL population from high density bin mapping using least square means at each environment and across two field experiments based on Best Linear Unbiased Prediction (BLUP) estimates.

**Figure 1.**
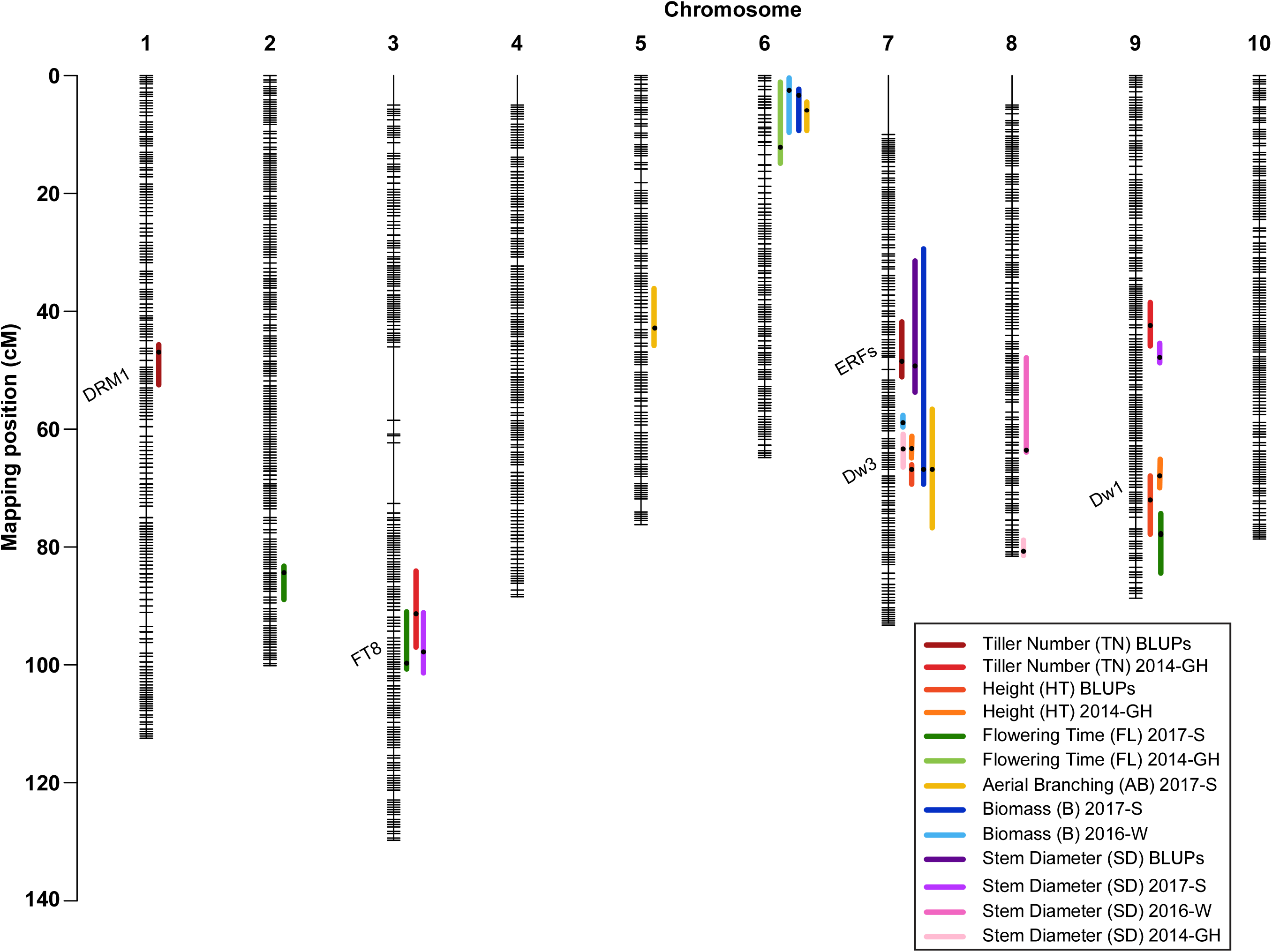
Genetic map containing QTL locations for six traits from 170 common RILs studied at both field locations. The map position in cM is shown on the y-axis. Horizontal lines on each chromosome represent a single bin marker while empty spaces represent regions containing heterozygous or duplicate markers, or lacking polymorphic markers. Vertical colored lines show the locations of QTL, with the color codes for each trait indicated in the inset.

#### Height

Across the three experiments, *S. propinquum* was taller (mean = 227 cm) than Tx7000 (mean = 96 cm), and the average plant height of the RIL population was 176 cm, ranging from 10 - 439 cm (**Table 4**). Plant height showed a fairly symmetrical distribution (skewness < 1) in the 2014-GH (**Supplemental Figure S1c**) and in the 2017-S experiments (**Supplemental Figure S4b**). In 2016-W, plant height was bi-modally distributed (**Supplemental Figure S4b**). Common QTL for height were detected on chromosomes 7 and 9 in the field trials. The QTL peaks for the BLUP summary analysis were located at 59.83 and 57.05 Mb, spanning 300 kb and 1.1 Mb intervals (qHT7.blup, qHT9.blup, respectively) (**Table 5 and Figure 1**). The BLUP summary explained 39.1 percent of the phenotypic variation. These QTL had an average negative additive effect of −17.3, indicating that Tx7000 alleles negatively influence height. In the greenhouse, the same QTL peaks for height were detected at 59.83 Mb (chromosome 7) and 56.68 Mb (chromosome 9), spanning 950 kb and 2.12 Mb, respectively. The two QTL explained at total of 28 percent of the phenotypic variance.

#### Stem diameter

Across the two field sites, *S. propinquum* had a smaller stem diameter (mean = 1.40 cm) than Tx7000 (mean = 1.85 cm). The average stem diameter across the RIL population was 1.73 cm, and ranged from 0.2 to 4.06 cm (**Table 4**). In the greenhouse, stem diameter was smaller than that observed in the field, with *S. propinquum* stems measuring an average of 0.92 cm, Tx7000 measuring an average of 1.14 cm, and a population mean of 0.93 cm (range of 0.1-2.5). Stem diameter displayed a normal distribution in all experimental trials (**Supplemental Figures S1c and S4c)**. QTL for stem diameter differed by environment. In the 2016-W experiment, QTL were detected on chromosomes 7 and 8 (qSD7.16W, peak at 53 Mb; qSD8.16W, peak at 60.43) (**Table 5 and Figure 1**). Together, these QTL explained 21.4 percent of the phenotypic variation with positive additive effects, indicating that Tx7000 alleles positively influenced stem diameter. In the 2017-S experiment, QTL on chromosomes 3, 7, and 9 were detected (qSD3.17S, peak at 62.64 Mb; qSD7.17S, peak at 4.78 Mb; and qSD.9.17S, peak at 48.32 Mb). These QTL together explained 36.5 percent of the phenotypic variation. The QTL on chromosomes 7 and 9 had positive additive effects (0.24) while the QTL on chromosome 3 had a negative effect (−0.13). The overlapping QTL on chromosome 7 between the 2016-W and 2017-S experiments ranged from 8.06 - 58.54 Mb and 3.22 - 58.69 Mb, respectively, but the location of the QTL peak differed between experiments (53 Mb in 2016-W and 4.78 Mb in 2017-S). The maximum QTL peaks from the BLUP summarized data were located on chromosomes 7 and 8 (qSD7.blup peak at 14.5 Mb, qSD8.blup peak at 59.4 Mb) and explained 23 percent of the phenotypic variation with additive effects indicating that Tx7000 alleles had a positive effect on stem diameter. With a similar effect, two QTL for stem diameter were detected on chromosomes 7 and 8 in the greenhouse experiment (qSD7.14GH peak at 59.9 Mb, qSD8.14GH peak at 62.4 Mb), and together they explained 21 percent of the phenotypic variation.

#### Biomass

The average plant biomasses of *S. propinquum* and Tx7000 were 1.14 kg and 0.07 kg in 2016-W, and 0.33 kg and 0.042 kg in 2017-S, respectively (**Table 4**). Biomass was not measured in the greenhouse trial. Biomass of the RIL population was positively skewed (skewness > 1, **Supplemental Figure S4d**), ranging from 0.01 to 2.79 kg plant^−1^ with a mean of 0.27 kg plant^−1^ across the two field environments. Common QTL for biomass were detected on chromosome 6 (qB6.16W, peak at 0.52; qB6.17S, peak at 0.87) and chromosome 7 (qB7.16W, peak at 58.63; qB7.17S, peak at 59.83) in the field experiments and explained ~18 percent of the phenotypic variance (**Table 5 and Figure 1**). In both field trials, the Tx7000 alleles had a negative effect on biomass; however, we observed a greater effect in 2016-W.

#### Flowering time

The number of days to flowering was recorded for the 2014-GH experiment and the 2017-S field trial. Tx7000 flowered after 50 days in the greenhouse, and 71 days in the 2017-S field study, while *S. propinquum* did not flower in either location (**Table 4**). For the plants that flowered (80% of the population, on average), flowering time was normally distributed and varied from 42 to 145 days with a mean of 84 days (**Table 4, Supplemental Figure S4e**). In the 2017-S field trial, three QTL for flowering time were identified on chromosomes 2 (qFL2.17S, peak at 73.31 Mb), 3 (qFL3.17S, peak at 63.32 Mb), and 9 (qFL9.17S, peak at 57.91 Mb), and explained 40.5 percent of the phenotypic variation (**Table 5 and Figure 1**). In the greenhouse, a flowering time QTL was detected on chromosome 6 (qFL6.14GH, peak at 39.97 Mb).

#### Aerial Branching

Aerial branching was measured only in the 2017-S field trial. *S. propinquum* produced no aerial branches, while Tx7000 produced an average of two aerial branches (**Table 4**). The frequency distribution for aerial branching was positively skewed, ranging from 0 to 49 with a mean of 6.3 (**Supplemental Figure S4f**). Three QTL for aerial branching were identified, one each on chromosomes 5 (qARB.17S. peak at 59.42), 6 (qARB.17S, peak at 2.28), and 7 (qARB.17S, peak at 59.74). These QTL explained 43.6 percent of the variation. The QTL on chromosomes 5 and 7 had average negative effects of −1.1 while the QTL on chromosome 6 had positive effects (**Table 5 and Figure 1**).

### Tiller bud transcriptome analysis

Differential expression analysis of tiller bud transcriptomes identified 6189 differentially expressed genes (DEGs) from the pairwise comparison between Tx7000 and *S. bicolor* subsp. *verticilliflorum* across a developmental series (10, 12 and 14 DAP) and within species between developmental stages (**Supplemental Table S4, Figure 2a**). K-means clustering of the 6189 genes identified seven co-expression clusters (**Figure 2b**). Clusters 4, 5 and 7 showed similar patterns of increased expression in Tx7000, with consistently lower expression in *S. bicolor* subsp. *verticilliflorum* across the same developmental stages, suggesting that these clusters contain genes involved in the acquisition of dormancy. In contrast, an increased expression level was observed in clusters 1, 2, 3 and 6 in *S. bicolor* subsp. *verticilliflorum,* suggesting that these clusters contain genes that may promote tiller bud elongation. Gene Ontology (GO) enrichment analysis identified several distinct regulatory processes associated with these clusters (**Figure 2c**). Cluster 2, which is upregulated in *S. bicolor* subsp. *verticilliflorum*, is enriched for genes involved in cellular and metabolic processes, while cluster 5, which is upregulated in Tx7000, is enriched for regulation of gene expression, and other regulatory functions. Cluster 7 was enriched for ADP, carbohydrate, nucleoside, and nucleic acid binding transcription factor activity involved directly or indirectly in regulating gene expression that controls bud growth or dormancy. The genes in clusters 1, 3, 4 and 6 were poorly annotated, so no significant GO terms were detected.

**Figure 2.**
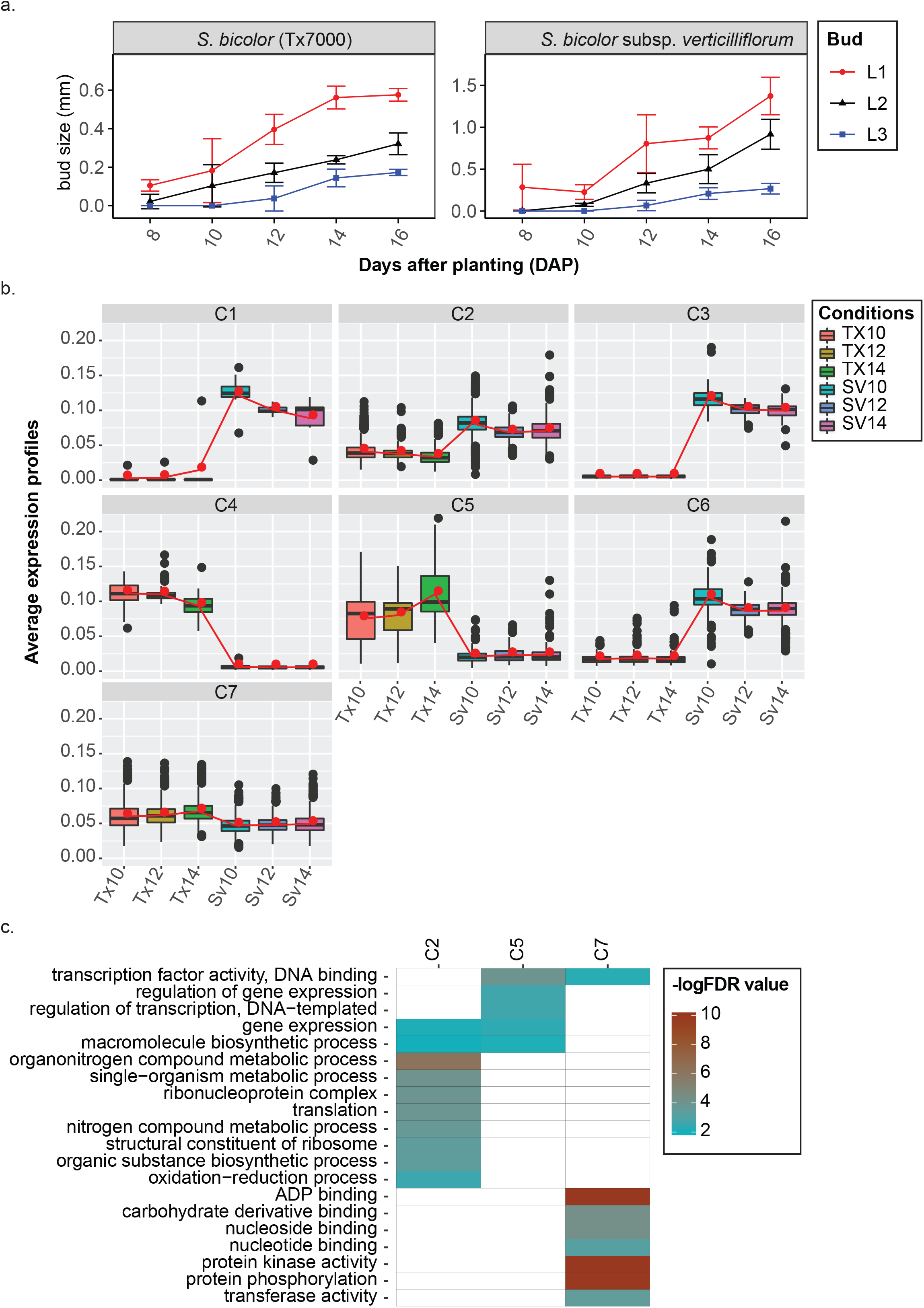
Transcriptional profile of tiller buds in *S. bicolor* Tx7000 and *S. bicolor* subsp. *verticilliflorum*. a) Length of tiller buds in the first (L1), second (L2), and third (L3) leaf axis of Tx7000 and *S. bicolor* subsp. *verticilliflorum* at 8, 10, 12, 14, and 16 days after planting (DAP). Buds from developmental stages at 10, 12, and 14 DAP were collected for RNA-seq transcriptome profiling. Bars for each stage represent the mean ± standard error of eight replicates. b) Seven co-expression clusters (C1 – C7) were identified from the 6189 genes that were differentially expressed across the developmental series in *S. bicolor* - Tx7000 and *S. bicolor* subsp. *verticilliflorum*. Clusters have been sorted such that those with similar mean vectors (as measured by the Euclidean distance) are plotted next to one another. Connected red lines correspond to the mean expression profiles for each cluster. The vertical bars define the upper and lower quartile, and the dots outside the bars indicate outliers. c) Over-represented GO terms in the co-expression clusters with false discovery rate (FDR) ≤ 0.01 (-logFDR ≥ 2) were considered as significantly enriched.

Of the 6189 DEGs, 409 were located within our tillering QTL windows. Among DEGs located in tillering QTL, 33% belong to co-expression clusters 2, 3, and 6 (upregulated in *S. bicolor* subsp. *verticilliflorum* tiller buds), and ~67% belong to co-expression clusters 4, 5, and 7 (upregulated in Tx7000 tiller buds). In the QTL on chromosome 1, there are 93 DEGs, of which 39 belong to clusters 2, 3 and 6 and 54 belong to clusters 4, 5, and 7. In the QTL on chromosome 7, the majority of DEGs (~75%) belong to clusters 4, 5, and 7, indicating increased expression in dormant buds. Two QTL were identified in the greenhouse experiment on chromosomes 3 and 9. Within these QTL there are 186 DEGs, of which ~34% show increased expression in *S. bicolor* subsp. *verticilliflorum* tiller buds, while the remaining 66% are upregulated in Tx7000 tiller buds.

We compared the genes in our co-expression clusters to a previous study that evaluated Phytochrome B (PhyB)-mediated regulation of tiller bud growth in sorghum (Kebrom and Mullet 2016). From differential expression analysis comparing wild type (tillering) to those of a phyB mutant (lacking tillers), the authors delineated several genes that appear to either promote or inhibit tiller bud initiation and elongation. Of the 78 DEGs presented in detail in Kebrom and Mullet, 48 were differentially expressed at 14 DAP, the time point at which Tx7000 buds acquired dormancy in our study (**Figure 2a**), and 39 of those displayed differential expression in the same direction (up or down-regulated). Seven of the 9 genes that displayed opposing expression patterns to those observed in Kebrom and Mullet are hormone related genes with differential expression during the early stages of bud development (up to 6 DAP), and therefore the discrepancy may be due to the differences in timing of the expression analysis (6 DAP vs 14 DAP).

## Discussion

In this study, we produced a high-density genetic map in sorghum composed of ~1920 bin markers spanning a genetic map length of ~875 cM constructed via low-coverage sequence-based genotyping. We used this population to identify QTL for tillering and other important sorghum traits, demonstrating its value as an important resource for the sorghum community. In comparison to previous studies that used low-density maps based on conventional marker systems such as restriction fragment length polymorphisms (RFLPs), amplified fragment length polymorphisms (AFLPs), and simple sequence repeats (SSRs) (Bowers et al. 2003; Hart et al. 2001; Kong et al. 2013; Lin et al. 1995), the genetic map produced here consisted of ~1.6 segregating markers per cM.

### Quality and accuracy of bin mapping

Plant height has been extensively studied in several sorghum QTL and genome wide association studies (Brown et al. 2008; Higgins et al. 2014; Morris et al. 2013), and all have pointed towards four major dwarfing genes (*Dw1*, *Dw2*, *Dw3* and *Dw4*) as the key contributors, either singly or in combinations, to the small stature of domesticated sorghum (Hart et al. 2001; Hilley et al. 2016; Hilley et al. 2017; Li et al. 2015; Lin et al. 1995; Quinby 1954). In our study, the two major QTL for plant height that were identified in all experiments overlapped with *Dw3* on chromosome 7 (qHT7.blup: 59.83 Mb; *Dw3*: 59.82 Mb) and *Dw1* on chromosome 9 (qHT9.blup: 57.05 Mb; *Dw1*: 57.04 Mb). Similarly, in the 2017 field trial, a flowering QTL was identified on chromosome 3 spanning a 4.0 Mb region (Chr03: 60.42 - 64.02 Mb) that contains a known member of the *Flowering locus T* (*FT*) family of flowering genes (Sobic.003G295300 at Chr03: 62.74 Mb; SbFT8; Wolabu and Tadege 2016), demonstrating the efficiency and accuracy of the bin mapping method used in this study. This result also indicates the degree to which different environments and/or different gene combinations can affect which contributing genes can be uncovered in a QTL study for even a highly penetrant trait.

### Segregation distortion

We identified several regions of highly distorted segregation, indicating gametic, zygotic, or inadvertent experimental selection during the formation of our RIL population. Two of the most distorted regions were on chromosomes 1 and 6, both showing biased transmission of *S. bicolor* alleles. On chromosome 1, the distal end was significantly enriched for *S. bicolor* alleles with an average marker ratio of 1:5 (SP:SB). This distorted region contains the maturity gene *Ma3* which is located near the distortion peak (phyB, Sobic.001G394400, Chr01: 68 Mb) (**Supplemental Figure 3**). The distorted region on chromosome 6 includes a large heterochromatic block (~50 Mb in size) containing the maturity gene *Ma6* (GDH7, Chr06: ~0.7) near the distortion peak, in addition to the SbFT-12 gene (Chr06: Sobic.006G047700 at 33.5Mb), and *Ma1* (SbPRR37 located at Chr06: 40.2 Mb). It is known that three maturity genes (*Ma1* [(SbPRR37, Chr06: 40.28 - 40.29 Mb]), *Ma2* [(Chr02: Sobic.002G302700 at 67.8 Mb]), and *Ma3* [(phyB, Sobic.001G394400, Chr01: 68.03 Mb]) are responsible for variation in flowering time among Milo genotypes (Casto et al. 2019; Quinby 1967). In addition, SbFT-12 is a floral suppressor during longer days (Cuevas et al. 2016), and is therefore expected to confer short-day length dependent flowering in *S. propinquum*-derived germplasm. Because the RIL population was advanced in a temperate latitude, it is likely that *S. bicolor* haplotypes were predominantly selected because of their ease of advancement, thereby causing segregation distortion of the regions related to photoperiod control of flowering.

Chromosomes 2 and 5 contained large segments demonstrating preferential transmission bias of *S. propinquum* alleles. The distorted region on chromosome 2 harbors the maturity gene, *Ma2*, near the peak of distortion (Chr02: Sobic.002G302700 at 67.8Mb, Casto et al. 2019). Interestingly, the combination of *Ma2* in its recessive state with *Ma1* is known to promote early flowering in long days in sorghum lines that are photoperiod sensitive (Quinby 1967). Therefore, it is likely that the *S. propinquum*-type lines that have lost day-length dependence were selected during RIL population development, in combination with *Ma1* on chromosome 6. If the cause of distortion relates to overall seedling or adult-plant vigor, a logical reason for inadvertent human selection during RIL population development, these *S. propinquum* regions may have value for *S. bicolor* improvement.

### QTL mapping

QTL for plant height were detected on chromosomes 7 and 9 in all experimental trials, indicative of strong genetic controls for this trait (**Figure 1**). On chromosome 7, the QTL overlaps with the sorghum dwarfing gene *Dw3* (Sobic.007G163800) and on chromosome 9 with *Dw1* (Sobic.009G229800). *Dw3* is a homologue of maize *Br2*, pearl millet *d2,* and *Arabidopsis* PGP1, and encodes a protein similar to ATP-binding cassette transporters of the multidrug resistant (MDR) class of P-glycoproteins (P-GPs) known to mediate polar auxin transport (Multani et al. 2003, Parvathaneni et al. 2013; Parvathaneni et al. 2019). *Dw1* encodes a putative membrane protein that is known to reduce cell proliferation activity in the internodes (Hilley et al. 2016; Yamaguchi et al. 2016). Further, the flowering QTL on chromosome 9 in the 2017-S field trail overlaps with *Dw1*. The colocalization of the flowering and height QTL on chromosome 9 mirrors that observed for the association that can be fractured between flowering time and plant height during mapping of sorghum conversion lines (Thurber et al. 2013), and therefore it is likely that the two QTL are controlled by different genes rather than an outcome of pleiotropy.

Unlike height, where common QTL were identified in all experimental trials, stem diameter demonstrated significant phenotypic plasticity in response to variable environmental conditions (**Figure 1**). Only one common QTL was identified on chromosome 7 across all experimental trials (qSD7.14GH, qSD7.16W, qSD7.17S); however, the QTL from the field experiments spanned a large chromosomal region, and their peaks were in different locations (qSD7.16W at 53 Mb; qSD7.17S at 4.78) (**Table 5**). The remaining QTL (qSD8.16W, qSD3.17S, qSD9.17S, and qSD8.14GH; **Table 5**) were unique to each environment. Unique QTL were also identified for tillering in the greenhouse experiment (qTN3.14GH, qTN9.14GH). There is a known non-causal negative relationship between stem diameter and tillering that is likely due to common hormonal control (Alam et al. 2014). Therefore, the significant variation in tillering observed among the experiments may explain the observed phenotypic plasticity and numerous unique QTL for stem diameter.

Similarly, unique QTL were identified for flowering time (**Figure 1**). The novel flowering time QTL qFL3.17S encompasses 433 annotated genes, including a gene that encodes a phosphatidylethanolamine-binding protein (PEBP) domain characteristic of members of the “FT” family of flowering genes (Sobic.003G295300; SbFT8; Chr03: 62.74 Mb, Wolabu and Tadege 2016). In response to short days, PEBP domain-containing genes promote flowering in photoperiod-sensitive sorghum genotypes (Wolabu and Tadege 2016). Wolabu and Tadege (2016) demonstrated accumulation of SbFT1, SbFT8, and SbFT10 transcripts in the leaf near the critical time of floral transition, suggesting that these three genes could be the sources of sorghum florigen, a flowering hormone. It is possible that SbFT8 and *Ma2* are working with other regulatory factors to affect the expression of *Ma1* during floral initiation (Casto et al. 2019). These results suggest the usefulness of this population in identifying genetic regulators of phenotypic plasticity to more fully delineate the genotype-phenotype relationship in diverse environmental conditions.

### Candidate genes for tillering in sorghum

To further evaluate the effectiveness of our population in identifying loci associated with important agronomic traits in sorghum, we combined QTL mapping with RNA-seq analysis during early tiller bud development. Additionally, we compared our results to those of Kebrom and Mullet (2016) in which the authors evaluated Phytochrome B (PhyB)-mediated regulation of tiller bud growth in sorghum. In the Kebrom and Mullet study, during the period from seed germination to six days after planting, the wild type and phyB mutant displayed similar patterns of bud formation and growth. During this period, genes related to cytokinins, gibberellic acid, and sugar transporters were differentially expressed (Kebrom and Mullet 2016). After 6 days, during the onset and development of bud dormancy, genes such as those encoding Dormancy Associated Protein 1 (DRM1, a well-known marker of bud dormancy), other dormancy related genes (e.g., NAC domain-containing proteins), and various transcription factors (WUSCHEL and Bellringer) were upregulated in dormant buds, while ACC oxidase, an ethylene responsive gene, and several early nodulin genes were downregulated.

Our results paralleled those of Kebrom and Mullet, particularly their findings during the onset and development of bud dormancy. There are 93 DEGs in the QTL on chromosome 1, of which 28 belong to cluster 2 and 49 belong to cluster 7, the most notable of which is DRM1 (Sobic.001G191200 at 16.9 Mb). DRM1 is upregulated ~2.3 fold in Tx7000 tiller buds and is likely the gene controlling this QTL. In the QTL on chromosome 7, the majority of DEGs (~75%) belongs to clusters 4, 5, and 7 and they include numerous transcription factors such as BTB-POZ, WRKY, and AP2/ERF. This QTL region also includes WUSCHEL (Sobic.007G087600, co-expression cluster 2), a homeobox transcription factor that is upregulated ~2.7 fold in *S. bicolor* subsp. *verticilliflorum* tiller buds and is known to be a positive regulator of tiller growth in rice (Wang et al. 2014). Other DEGs from co-expression cluster 2 that lie within QTL identified in at least one of the environments include genes involved in the generation of cytokinins (CKs), which are known to promote bud outgrowth (Müller et al. 2015), and abscisic acid (ABA) related genes that are known to repress bud outgrowth (Chatfield et al. 2000; Reddy et al. 2013). Additionally, it should be noted that there are several DEGs of unknown function that belong to co-expression clusters 2 and 6 and are located within the two QTL from the field trials. These DEGs are upregulated in *S. bicolor* subsp. *verticilliflorum* tiller buds, and in several cases no expression was detected in Tx7000 tiller buds. These results demonstrate the power of combining QTL mapping with RNA-seq for the discovery of novel genes that control a phenotype of interest.

Several other genes show similar patterns of expression but are located in QTL identified from only one of the experimental environments, suggesting that they play a role in the environmental plasticity of tiller development. These genes include regulators of abscisic acid levels, transcription factors, and gibberellin-regulated family proteins, all known to be involved in promoting bud dormancy. Interestingly, two important transcription factor genes, *tb1* (SbTb1, Sobic.001G121600) and *gt1* (SbGt1, Sobic.001G468400), known to regulate axillary branching in maize (Dong et al. 2019), were not identified as DEGs in this study; however, there was a 1.5-fold increase in SbGt1 gene expression in dormant buds of Tx7000 relative to elongating buds of *S. bicolor* subsp. *verticilliflorum*, although it was not judged statistically significant between time points. Similarly, Kebrom and Mullet did not detect differential expression of *tb1* using RNA-seq, although qRT-PCR validation experiments detected a 2-fold increased expression of *tb1* in the PhyB mutant.

## Conclusions

As in all QTL studies with only two parents, finite progeny population sizes, and testing across a limited number of environments, it is not expected that all contributing genetic variation for a trait will be discovered. Therefore, it was impressive in our analysis that several QTL were found in two or more experiments across widely different environments and dates. For example, QTL for plant height were detected on chromosomes 7 and 9 in the greenhouse and in both experimental field trials. It was also interesting that many of the 29 QTL that we found for the most penetrant traits were associated with already-known genes. In all of our field and greenhouse experiments, and for all traits analyzed, transgressive segregation was the rule, with progeny often exhibiting several fold greater variation for a trait than seen when comparing the parents. This is a routine observation in plant genetics when crosses between domesticated and wild species are analyzed (Lippman and Tanksley 2001; Cong et al. 2002; Doust et al. 2004; Tanksley 2004; Mauro-Herrera and Doust 2016; Singh et al. 2018), but it does emphasize the tremendous potential value of this population for future studies of domestication and crop improvement. Most exciting, many QTL were found in regions that lack well-known genes that could account for the detected variation. The combination of QTL mapping with RNA-seq is a powerful approach for identifying such candidate genes. This now-available RIL population and these results provide important targets for future study and improvement of *Sorghum bicolor*.

## Acknowledgments

The authors wish to thank Michael Boyd, Yisel Carrillo, Matt Estep, Molly Haddox, Alex Harris, John Hodge, Aye Htun, Jacob A. Kelly, Melissa Lehrer, Qing Li, Jonathan Markham, Torrey Nielsen, Ben Purintun, Dhanushya Ramachandran, and Maddie Squire for their assistance in data generation, collection, and analysis. We acknowledge the West Virginia University Evansdale Greenhouse for supplying space. We acknowledge the help of Josh Massey and the field staff at the Cimarron Valley Research Station, Perkins, OK, and of the field staff at the University of Georgia’s Iron Horse Farm, Watkinsville, GA for providing and preparing the field space. We would also like to acknowledge the WVU Genomics Core Facility, Morgantown, WV for support provided to help make this publication possible and CTSI Grant #U54 GM104942 which in turn provides financial support to the WVU Core Facility. Additional funding information: The BioEnergy Science Center (Department of Energy, United States) (to JLB), The Center for Bioenergy Innovation (Award DE-AC05-000R22725) (to JLB and KMD), and The National Science Foundation IOS-1339332 (to AND, JSH, CW).

## Supplementary Tables and Figures

**Supplemental Table S1. Analysis of variance summary.** Sum of square values (type III) from combined analysis of variance for traits measured in RILs in Watkinsville, GA (2016) and Stillwater, OK (2017) for tiller number, height, and stem diameter.

**Supplemental Table S2.** Summary statistics for RNA-seq experiment

**Supplemental Table S3.** RNA-seq count matrix used in the transcriptome analysis

**Supplemental Table S4.** Differentially expressed genes in tiller buds identified within and between species comparisons across 10, 12, and 14 DAP. This gene set was subjected to co-seq cluster analysis. The co-seq cluster ID for each gene is shown in column B. The average expression profile calculated in the co-seq analysis (columns C-H) and the respective RPKM values (columns N-S) are shown.

**Supplemental Figure S1.** Pearson correlation matrix for each phenotype measured in all three experiments. a) 2016-W field trial, b) 2017-S field trail, c) 2014-GH greenhouse experiments. The distribution of each phenotype is shown on the diagonal. Below the diagonal is the bivariate scatterplot with a line of best fit. Above each diagonal is the Pearson correlation coefficient and associated significance level. Significance levels, indicated by asterisk(s), are associated with a p-value threshold (p < 0.001 (***); p < 0.01 (**); p < 0.05 (*). HT = Height; TN = Tiller Number; SD = Stem Diameter; FL = Flowering Time; ARB = Aerial Branching.

**Supplemental Figure S2. Recombination bin-map of the RIL population.** The bin-map consists of 3652 bin markers for 170 recombinant inbred lines that were common to both field experiments. Physical positions are based on the *S. bicolor v*3.1 genome sequence. Red: *S. propinquum* homozygote; Green: *S. bicolor* Tx7000 homozygote; Blue: heterozygous.

**Supplemental Figure S3. Segregation distortion.** The x-axis indicates the physical position of bin markers on the chromosomes, and the y-axis shows the negative logarithm of p-values with a base of 10. The map locations of some of the known important flowering time genes in sorghum are depicted.

**Supplemental Figure S4.** Frequency distribution for phenotypes used in QTL analysis. Histograms illustrate the frequency of traits in the 2016 Watkinsville, GA and the 2017, Stillwater, OK field experiments. (A) Tiller number, (B) Plant height, (C) Stem diameter, (D) Biomass, (E) Flowering time, and (F) Aerial branches.

