## Supplementary figures and images for "Integration of high-density genetic mapping with transcriptome analysis uncovers numerous agronomic QTL and reveals candidate genes for the control of tillering in sorghum"

### SupplementalFigureS1

a.

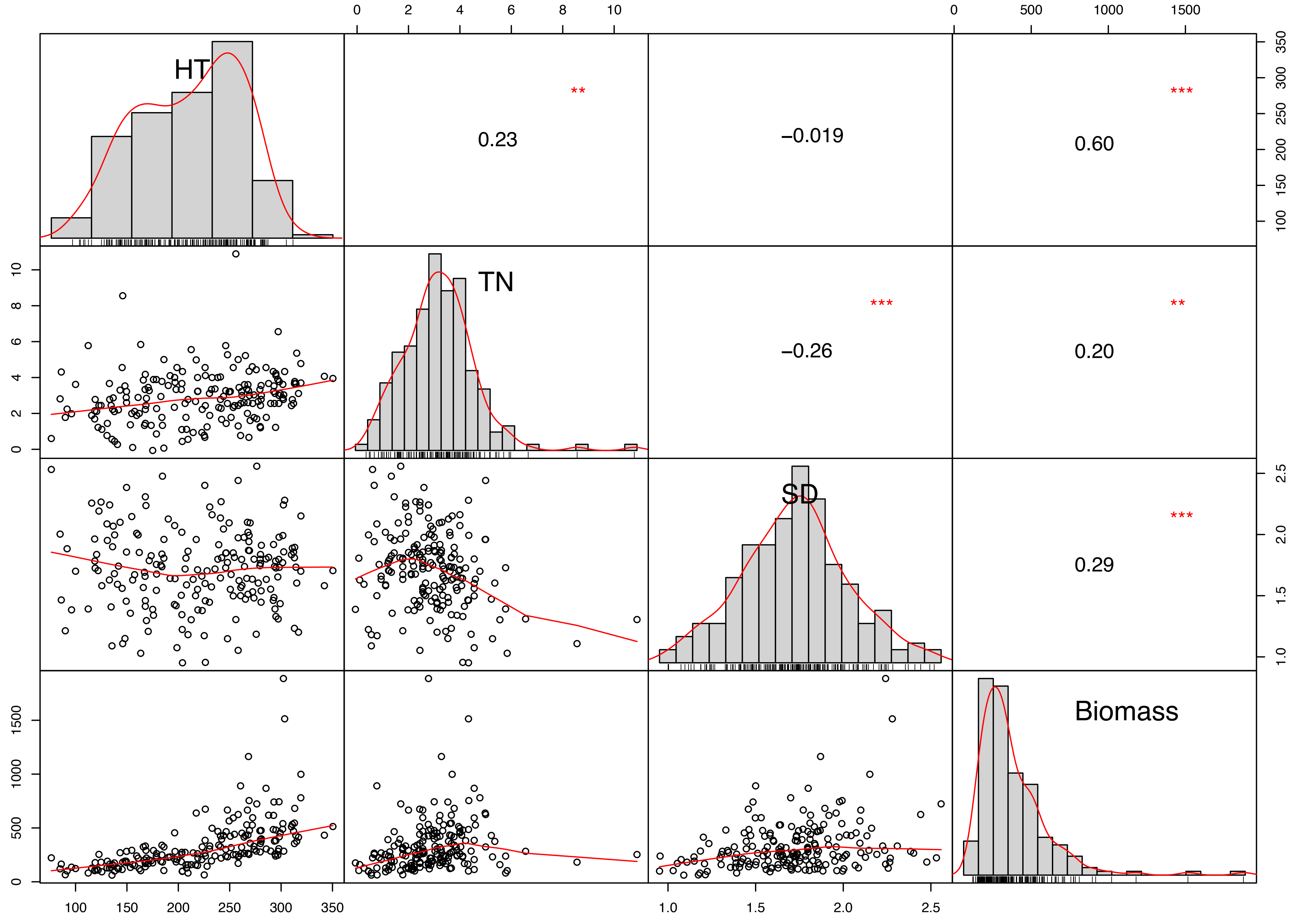

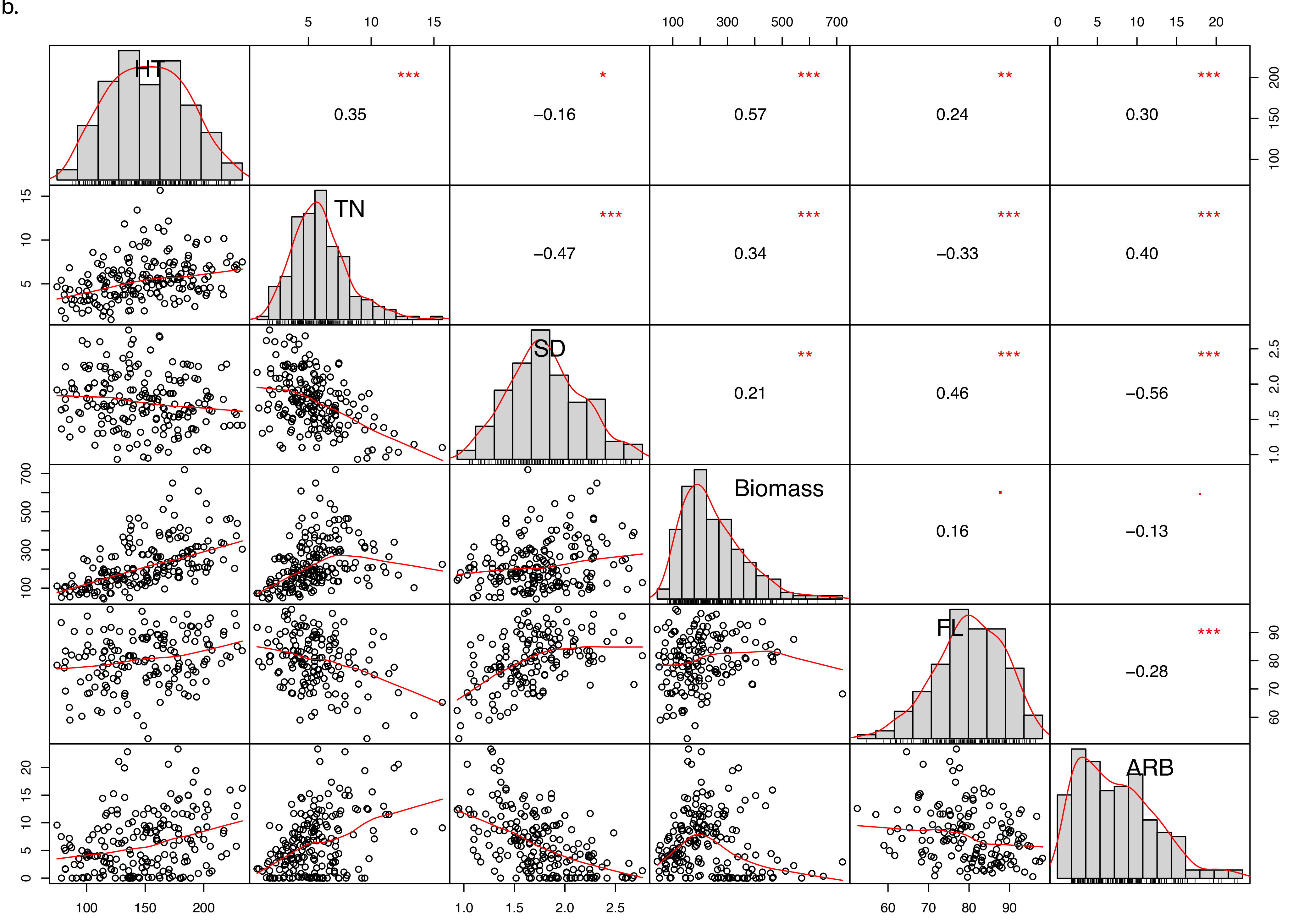

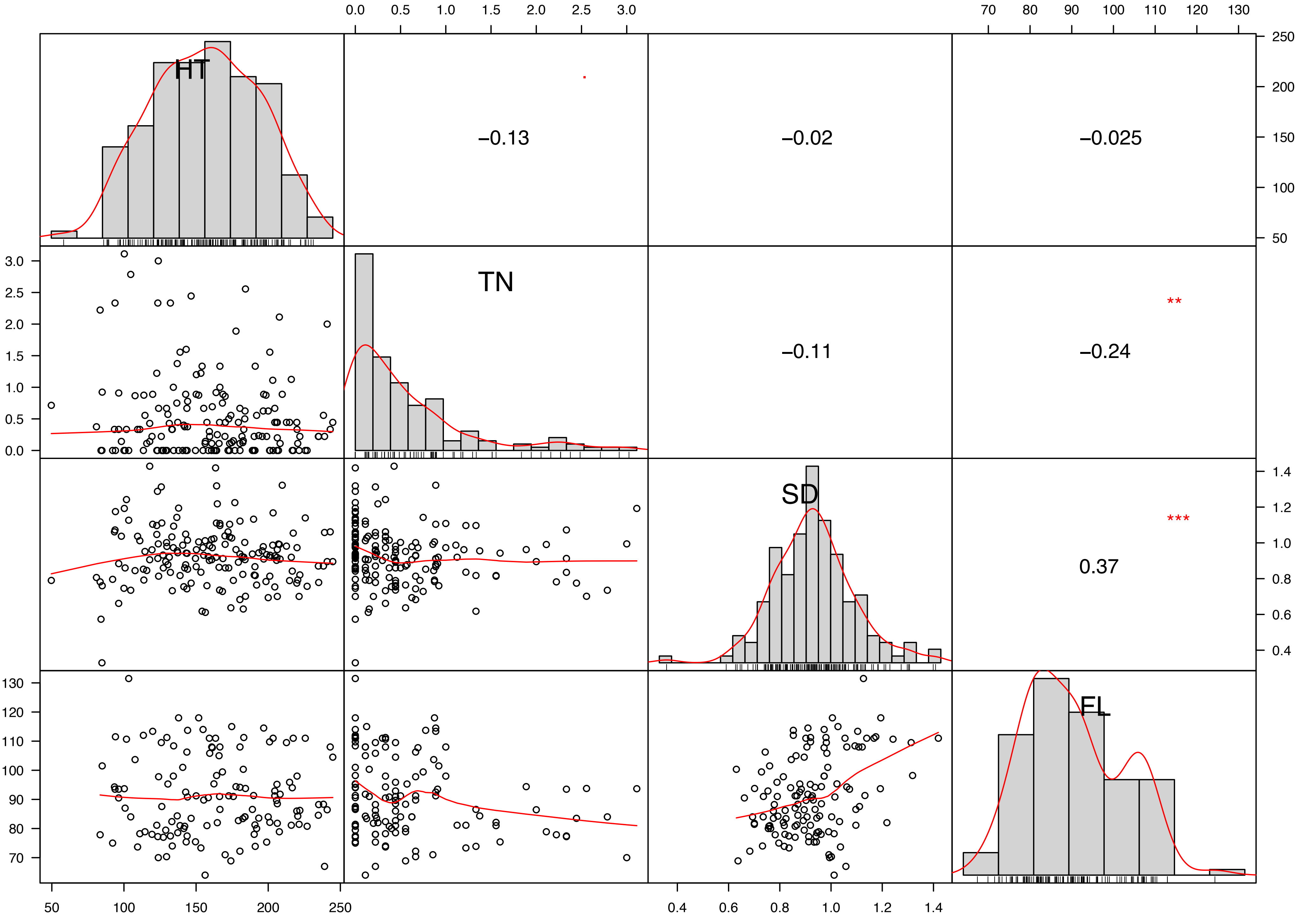

### SupplementalFigureS2

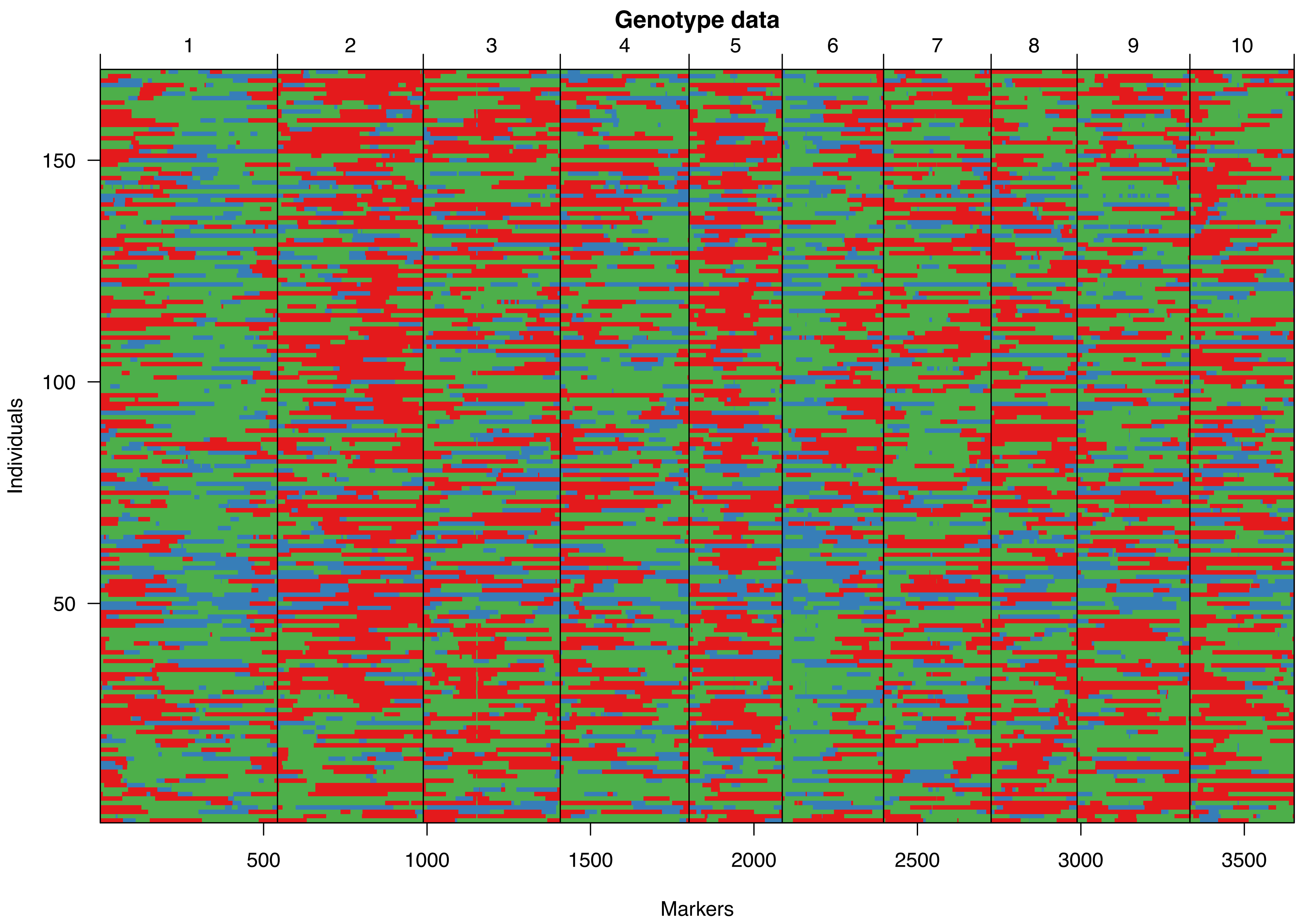

### SupplementalFigureS3

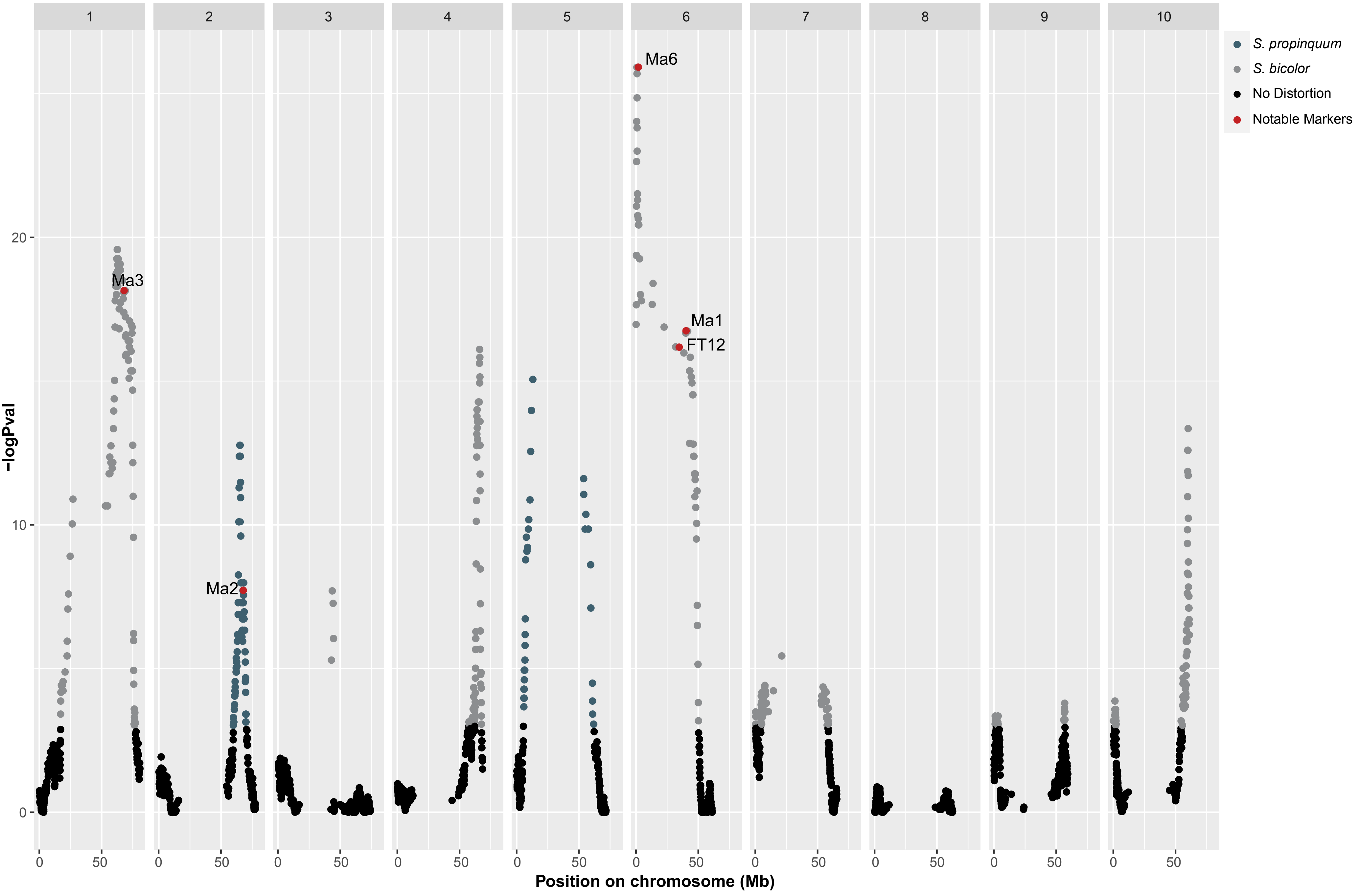

### SupplementalFigureS4

**Supplemental Figure S4. Frequency distribution for phenotypes used in QTL analysis.**

a.

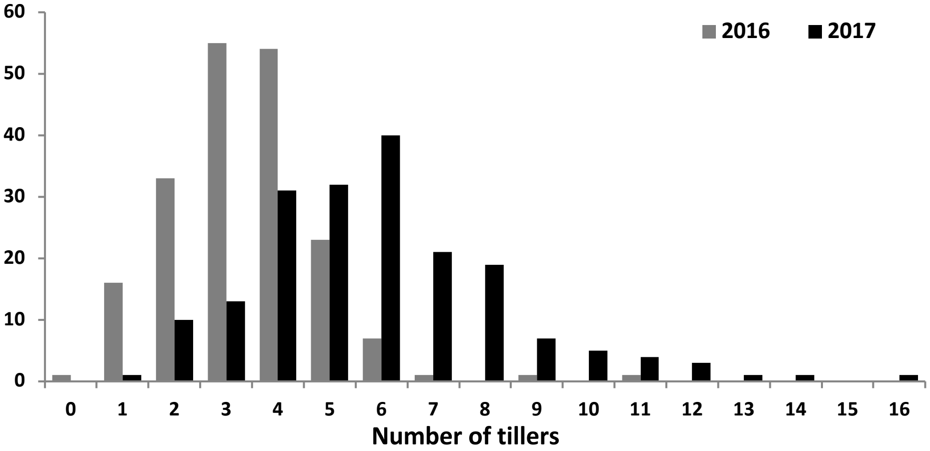


b.

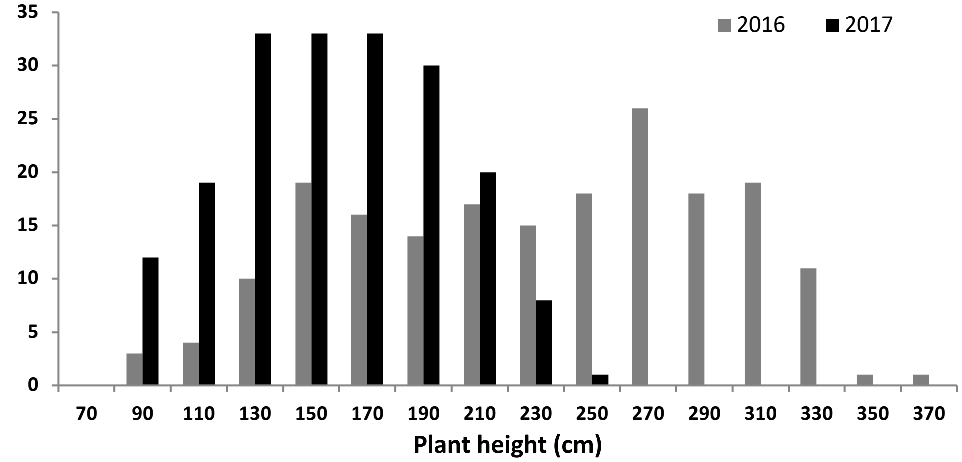

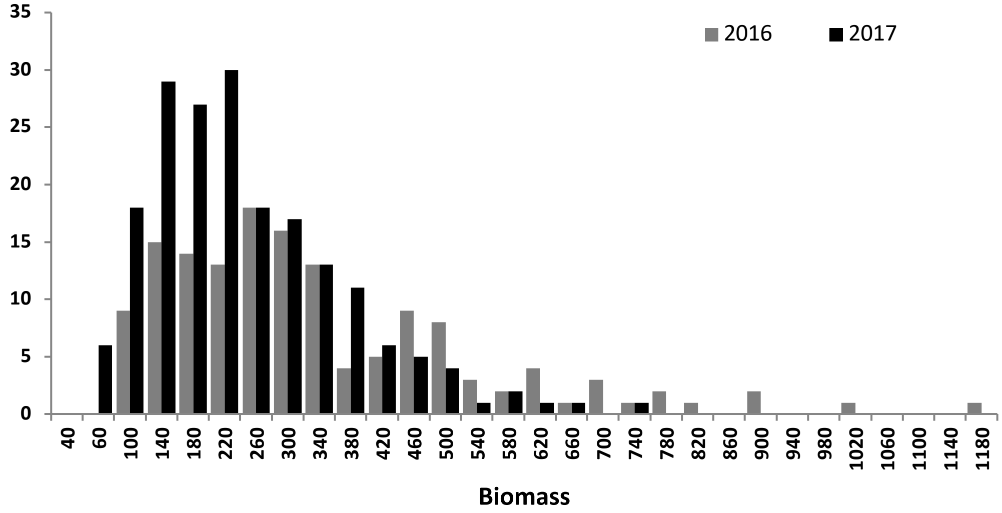

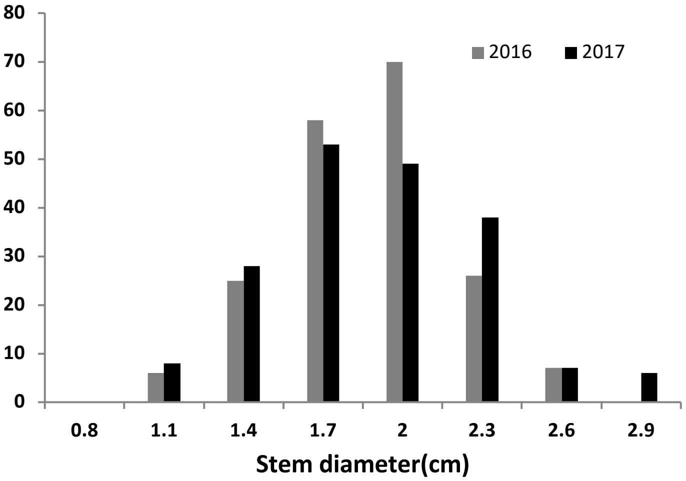

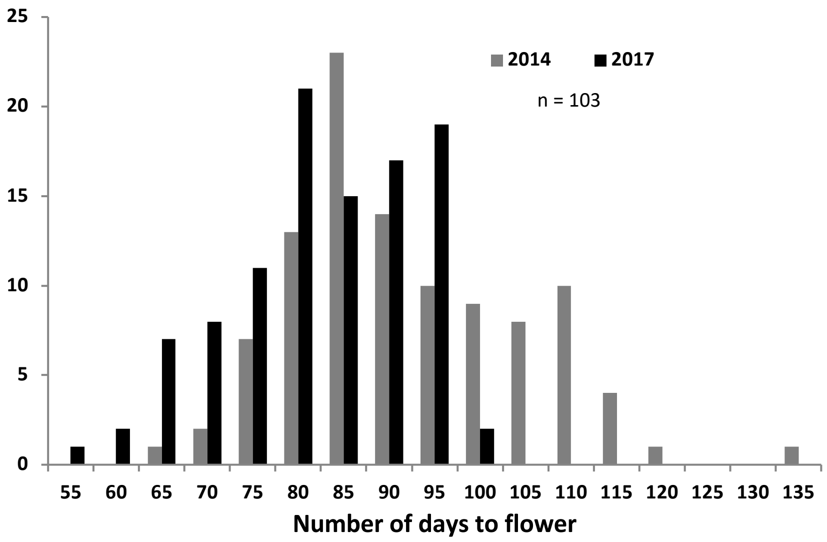

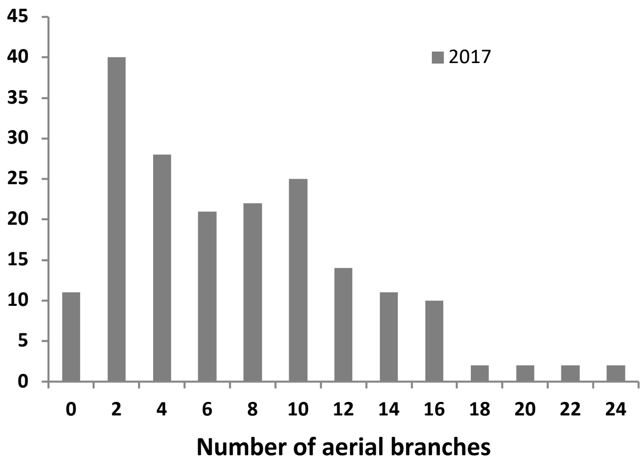
